# Photoperiod modulates mu-opioid receptor availability in brown adipose tissue

**DOI:** 10.1101/2022.04.08.487600

**Authors:** Lihua Sun, Richard Aarnio, Erika Atencio Herre, Salli Kärnä, Senthil Palani, Helena Virtanen, Heidi Liljenbäck, Jenni Virta, Aake Honkaniemi, Vesa Oikonen, Chunlei Han, Sanna Laurila, Marco Bucci, Semi Helin, Emrah Yatkin, Lauri Nummenmaa, Pirjo Nuutila, Jing Tang, Anne Roivainen

**Affiliations:** Turku PET Centre, University of Turku, FI-20520 Turku, Finland; Turku PET Centre, Turku University Hospital, FI-20520 Turku, Finland; Department of Nuclear Medicine, Huashan Hospital, Fudan University, Shanghai, China; Turku Center for Disease Modelling, University of Turku, FI-20520 Turku, Finland; Heart Center, Turku University Hospital, FI-20520 Turku, Finland; Central Animal Laboratory, University of Turku, FI-20520 Turku, Finland; Department of Psychology, University of Turku, FI-20520 Turku, Finland; Department of Endocrinology, Turku University Hospital, FI-20520 Turku, Finland; InFLAMES Research Flagship Center, University of Turku, FI-20520 Turku, Finland; Research Program in Systems Oncology, Faculty of Medicine, University of Helsinki, FI-00014 Helsinki, Finland

**Keywords:** mu-opioid receptor, brown adipose tissue, photoperiod, innervation, positron emission tomography, carfentanil

## Abstract

Photoperiod drives metabolic activity of brown adipose tissue (BAT), and affects food intake and weight gain in mammals. Sympathetic innervation in BAT controls thermogenesis and facilitates physiological adaption to seasons, but the exact mechanism remains elusive. Previous studies show that the central opioid signaling tunes BAT heating and the brain mu-opioid receptor (MOR) levels have seasonal patterns. It is hence intriguing to know whether the peripheral MOR signaling shows seasonal variation. Here, we examined the effect of photoperiod on BAT MOR availability using [^11^C]carfentanil positron emission topography (PET). Adult rats (n = 9) were repeatedly imaged under changing photoperiods which simulates the local seasons. Long photoperiod downregulated MOR availability in BAT, while MOR availability in the muscles was unaffected. We confirmed the expression of MOR in BAT and muscle using immunofluorescence imaging. We conclude that photoperiod causally affects MOR availability in BAT, and sympathetic innervation of BAT may influence thermogenesis via the peripheral MOR system.

**Significance of the study:** Photoperiod impacts the metabolic activity of brown adipose tissue (BAT) with the exact mechanism still unclear. The current study shows that photoperiod causally affects the mu-opioid receptor (MOR) levels in BAT, with longer photoperiod leading to lower MOR availability. This possibly indicates down-regulated innervation during bright seasons. Immunofluorescence staining data reveal expression of MOR in both brain and peripheral tissues, drawing attention to the under-investigated peripheral MOR system. Also, the study highlights the feasibility of [^11^C]carfentanil PET in studying the peripheral MOR signaling.

## Introduction

Brown adipose tissue (BAT) is known for rapid production of heat via non-shivering thermogenesis, and the capacity of this function demonstrates seasonal rhythms. Environmental factors such as photoperiod drive the BAT metabolic activity. Short photoperiod remarkably stimulates BAT growth and thermogenesis in Syrian hamsters, along with increased food intake and weight gain (McElroy and Wade, 1986). This effect of photoperiod on BAT thermogenesis seems to be independent from the parallel effect of environmental temperature variation (Wiesinger et al., 1989; Au-Yong et al., 2009), suggesting a distinctive physiological pathway in developing this adapted seasonal rhythm. Exact neurochemical effectors underlying the seasonal patterns of BAT function remains elusive. In humans, *in vivo* positron emission tomography (PET) imaging evidence also reveals increased BAT glucose uptake in winter (Ouellet et al., 2011) and meanwhile, the human feeding patterns show increased caloric intake from fats in the fall and winter (Shahar et al., 1999; Ma et al., 2006). Recent studies suggest that the role of BAT may be far beyond the traditionally known thermogenesis, by conceptualizing a functional BAT-brain axis that tunes neurometabolic control and feeding behavior (Li et al., 2018; Laurila et al., 2021; Virtanen and Nuutila, 2021). Interestingly, among these multiple perspectives of BAT functions, photoperiod seems to deliver a fine-tuning effect eventually shaping the seasonal patterns of metabolism and food reward related behavior in both animals and humans.

### Central and peripheral opioid systems

The endogenous opioid system regulates pain, stress and emotions (Darcq and Kieffer, 2018), and especially the central mu-opioid receptor (MOR) signaling is also a potent modulator of feeding (Karlsson et al., 2015; Tuulari et al., 2017; Nummenmaa et al., 2018). The role of central MOR system in feeding behavior is thought to reside in its contribution of hedonic or “liking” responses in the brain (Berridge, 2009). We have recently shown that photoperiod drives seasonal patterns of central MOR availability and affects weight gain in rats (Sun et al., 2021). Compared to central MOR signaling, understanding of peripheral opioid functions has been mainly based on studies of pain (Vadivelu et al., 2011), as exemplified by the local analgesic effect of opioid peptides (Stein et al., 1993). In peripheral nerves, opioid receptors are synthesized at the dorsal root ganglion and transported intra-axonally to terminals (Fields et al., 1980; Young et al., 1980), and opioid receptors have also been found in the peripheral sensory nerve terminals and immune cells (Hassan et al., 1993; Zhou et al., 1998). Tissue injury augments peripheral opioid analgesia along with increased MOR density (Zhou et al., 1998; Kraus et al., 2001; Stein et al., 2001), suggesting an engaged role of MOR signaling in immune system and inflammation. However, general knowledge on the peripheral opioid receptor functions, for example their roles in feeding (Delgado-Aros et al., 2003), is still very limited.

### The opioid system and BAT metabolism

BAT has high number of sympathetic nerves that controls thermogenesis (Kuruvilla, 2019). Neuronal release of noradrenaline activates BAT adrenergic receptors thus to stimulate biochemical reactions in mitochondria and thermogenesis (Bartness et al., 2010). On the other hand, BAT synthesizes protein calsyntenin 3β to promote the growth of projections from neurons (Zeng et al., 2019), thus to strengthen innervations. The role of opioid signaling as a modulator of BAT thermogenesis is supported by data showing that intravenous and intracerebroventricular administration of fentanyl enhances BAT sympathetic nerve activity and thermogenesis (Cao and Morrison, 2005). There are no reports on the role of peripheral MOR signaling and BAT activation. However, inflammatory response in BAT has been associated with reduced capability of thermogenesis (Omran and Christian, 2020) and inflammation is meanwhile related to augmented MOR signaling (Stein et al., 1993, 2001) indirectly pointing to possible roles of peripheral MOR in BAT activity. Interestingly, both mammalian immunity and BAT thermogenesis demonstrates seasonal patterns (McElroy and Wade, 1986; Stevenson and Prendergast, 2015).

### The current study

In the current study, we tested whether the peripheral MOR signaling is involved in photoperiod-modulated innervation of BAT activity. Rats (n = 9) were kept under varying photoperiods simulating local seasonal cycles, with each rat imaged repeatedly using [^11^C]carfentanil PET. Using this dataset, we have previously reported that photoperiod shapes *in vivo* brain MOR availability (Sun et al., 2021). Meanwhile, in the same study we have also reported that seasonal cycling of photoperiod elevates stress hormone and slows down weight gain. Here, we further studied the effect of photoperiod on BAT MOR availability. We hypothesize that photoperiod tunes innervations of BAT functions via the MOR signaling pathway.

## Methods

### Animal handling and seasonal simulation

Eighteen adult Sprague-Dawley rats (Envigo, Horst, Netherlands; age > 90 days; 11 males, 7 females) were housed under controlled laboratory conditions in open top cages with free access to water and CRM-E diet (SDS, UK). Rats were caged in groups of two or three same-sex rats. The study included an experimental group (9 males, 5 females), for which the in-house light with variable ON/OFF duration was programmed; cycling photoperiods simulate the local seasonal change of daylength with a speeded cycle completed in three months. We also involved a control group (2 males, 2 females), which was kept in a different room with constant daylength cycle (12h ON/ 12h OFF), with all other conditions the same with the experimental group. The control group was used for addressing the potential effects of ageing on MOR levels that occur in the absence of variable daylength; the main statistical analyses pertaining to MOR were also run separately in the experimental group only. The same type of LED lighting was used for control groups and the experimental groups. The study was in accordance with the EU Directive 2010/63/EU on the protection of animals used for scientific purposes based on the 3R principles, and all the procedures and protocols were approved by the National Project Authorisation Board of Finland (Licence numbers 3116/04.10.07/2017 and 8648/2020).

### PET imaging and processing

Twelve out of eighteen rats (experimental group: 6 males, 3 females; control group: 2 males, 1 female) were studied with 60 minutes dynamic [^11^C]carfentanil PET imaging for 3-4 times under isoflurane anaesthesia. The radiotracer was divided by three rats at each scanning day (2 males, 1 female), and via using two different scanners to maximize the data collection per batch. Due to larger body size, Inveon Multimodality PET/CT (Siemens Medical Solutions, Knoxville, TN, USA) was used for imaging the male rats (two rats at a time), whereas Molecubes PET/CT (Gent, Belgium) was used for the female rats. In total, there were 42 PET/CT scans. Rats were weighed on the scanning day. For the Inveon Multimodality PET/CT scanner, with relatively lower resolution and sensitivity, the aimed injected radioactivity dose of [^11^C]carfentanil was 5 MBq, with actual injected dose of 4.69 ± 0.60 MBq corresponding to weight-based mass dose of 29.28 ± 20.53 ng/kg. For Molecubes PET/CT scanner, the aimed dose was 1 MBq, with actual injected dose of 1.14 ± 0.13 MBq corresponding to 13.92 ± 11.74 ng/kg. While there may be receptor saturations, the saturation is similar in all conditions and thus does not affect the cross-condition comparisons. Accordingly, the effects of sex and dosage cannot be separated from scanner type and dosage dependent effects.

Dynamic PET images were analysed using Carimas software (version 2.10.3.0) developed at the Turku PET Centre, Finland. The PET data sets were reconstructed into 20 time frames using three-dimensional ordered subset expectation maximization (OSEM3D) algorithm: 6 × 0.5 minutes, 3 × 1 minutes, 4 × 3 minutes, and 7 × 6 minutes. Regions of interests (ROIs) were defined independently by two researchers (HV and EAH) who were blinded to the experimental conditions.

### Immunofluorescence staining

Brains, BAT, muscle, stomach and white adipose tissue from healthy male Sprague-Dawley rats (n=3, kept under a regular 12h/12h dark-light conditions and not involved in the PET imaging study) were collected on ice, frozen and cut into 10 μm cryosections. Brains were cut into transverse sections.

Tissue sections were fixed in 10% formalin for 10 minutes and permeabilized in 0.1% Triton X-100 in phosphate-buffered saline (PBS) for 5 minutes, and washed between and after the incubations with PBS. For blocking of unspecific staining, sections were incubated in 5% normal goat serum (NGS, Vector laboratories) in PBS for 1 hour at room temperature. Then, sections were incubated with anti-OPRM1 (1:100 dilution with 2.5% NGS in PBS, #MAB8629, Bio-Techne, Minneapolis, Minnesota, USA) for overnight at +4 C. After PBS washes, sections were incubated with AlexaFluor 488-conjugated anti-rabbit secondary antibody (1:500 dilution with 2.5% NGS in PBS, Invitrogen, Waltham, Massachusetts, USA) for 1 hour at room temperature, followed by staining with 4’,6-diamidino-2-phenylindole (DAPI, 1:10,000 dilution with milliQ water, Sigma-Aldrich) for 10 minutes. Finally, the sections were mounted with Prolong Gold Antifade mounting media (#P369309, Thermo Fisher Scientific). Sections with omitted primary antibody were used to validate the specificity of staining. Images were obtained with 3i Spinning Disk confocal microscope using Slidebook software (Intelligent Imaging Innovations) and processed with ImageJ software.

### Radiometabolite analysis of [^11^C]carfentanil

Four female and three male rats (not involved in the PET imaging study) were anesthetized using isoflurane and injected intravenously with [^11^C]carfentanil (injected dose 39.5 ± 17.5 MBq) in order to assess *in vivo* stability of the tracer in blood plasma and different organs.

Blood samples (0.2-0.6 mL) drawn at 5 and 10 minutes after injection were collected during deep anaesthesia from the tail. At 20 minutes post injection, blood was collected by cardiac puncture and the rats were euthanized using cervical dislocation. The blood samples were collected into heparinized gel tubes (Microtainer, Becton, Dickinson and Company, Franklin Lakes, NJ, USA) and centrifuged with 14,400 *×g* for 90 seconds. The separated plasma was mixed with ice-cold acetonitrile (plasma:acetonitrile 2:3 (*v*/*v*)) to precipitate the plasma proteins and centrifuged with 14,400 *×g* for 90 seconds. The supernatants were analysed using the high-performance liquid chromatography coupled with a radiodetector (radio-HPLC: Merck Hitachi L-7100 gradient pump system with Radiomatic 150 TR, Packard, USA). The brain, BAT, liver, stomach, muscle and duodenum were dissected and homogenized with 1:1 (*v*/*v*) methanol:water using an electric homogenizer (Ultra-Turrax T8, IKA, Staufen, Germany). The homogenized solutions were filtered (Amicon Ultra-4, 50k, Merck KGaA, Darmstadt, Germany) to remove particle residuals and proteins from clogging the HPLC column, and then injected to the radio-HPLC. In the radio-HPLC, a μBondapak C18 column (7.8 × 300 mm, 10 μm; Waters, Milford, MA, USA) with a gradient method with 50 mM phosphoric acid and acetonitrile was used, and the results were analyzed as previously described (Hirvonen et al., 2009).

### Modelling of the MOR availability

Standardized uptake values (SUVs) were calculated by dividing tissue radioactivity concentration (MBq/ml) with injected radioactivity dose per animal weight (MBq/g). One approach used to estimate the MOR availability was to calculate the SUV ratios between areas under the time-activity curves (TACs) of BAT and muscle (i.e. reference tissue). Alternatively, areas under the TACs can also be an approximate index of the MOR availability. Based on our radiometabolite analysis data, the first 5 minutes (due to confounding perfusion effects at the peripheral region) and the last 40 minutes (due to radiotracer decomposition) of the TACs were excluded in these modelling.

### Statistical analysis

Data of PET repeated measures were analyzed using a linear mixed-effects model with varying intercepts for each rat and fixed-effect factors such as age, photoperiod and scanner type. R statistical software (version 4.0.3) with the lme4 package was used.

## Results

### Peripheral MOR expression and [^11^C]carfentanil PET imaging

Immunofluorescence staining showed the most prominent expression of MOR in BAT and stomach (**Fig. 1A**). Relatively lower amount of MOR was expressed in muscle, and almost no expression was found in white adipose tissue. In line with the literature, MOR had higher expression in the neocortex compared to cerebellum in the brain (**Fig. 1B**).

**Figure 1.**
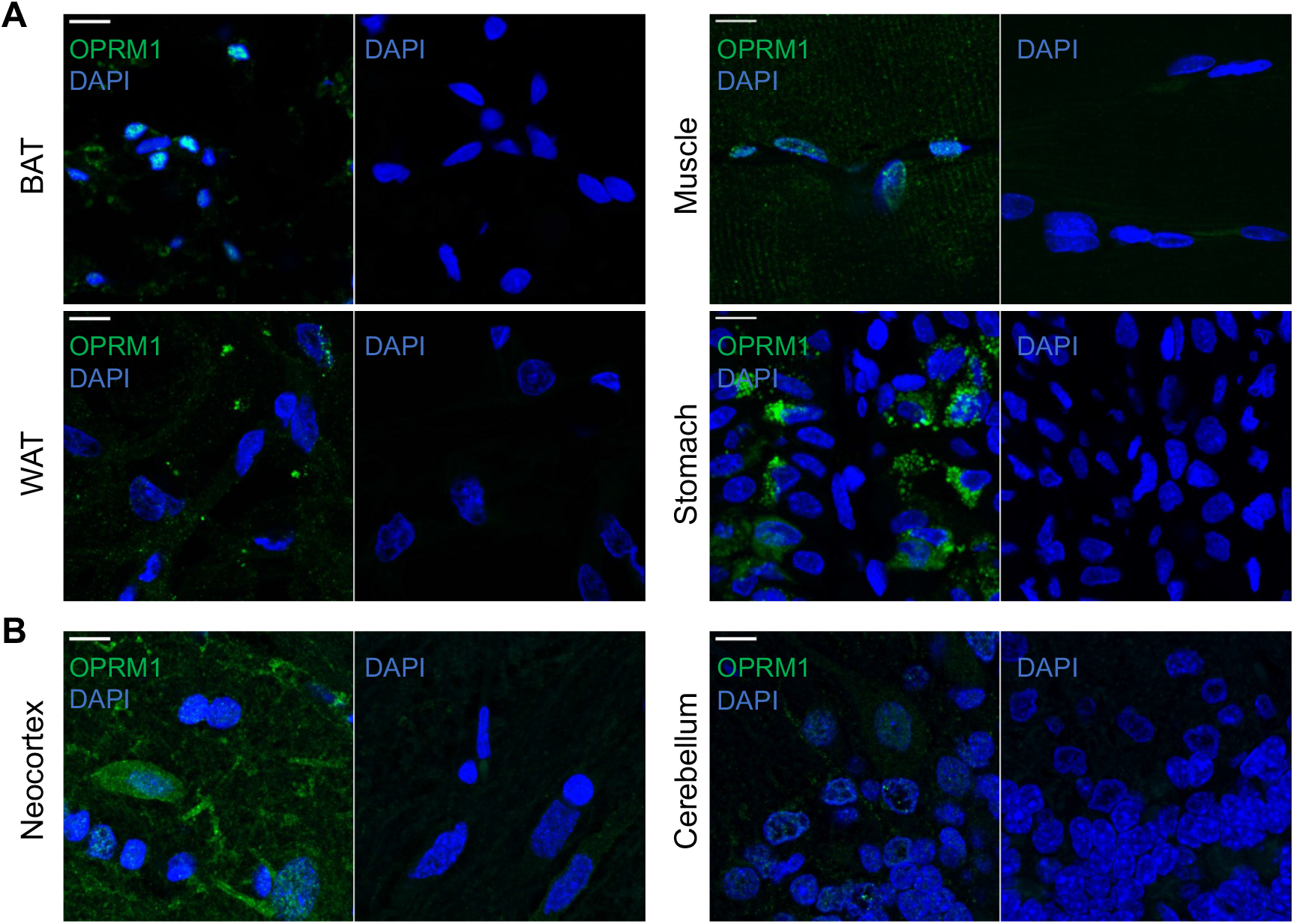
Representative immunofluorescence confocal microscopy images with anti-MOR antibody (OPMR1, green) reveals positive staining in peripheral (**A**) and brain tissues (**B**) in the left panels. Nuclei were counterstained with DAPI (4-6-diamidino-2-462 phenylindole, blue). Omission of anti-MOR antibody resulted in no staining in the right panels. BAT = brown adipose tissue; WAT = white adipose tissue. Scale bar = 10 μm.

In the analysis of PET images, we firstly calculated the standardized uptake values (SUVs) in the brain and five peripheral tissues including BAT, muscle, stomach, liver and duodenum (**Fig. 2A**), shown as the TACs from dynamic PET images (**Fig. 2B**). SUVs were the highest in stomach, liver and duodenum, while relatively lower in BAT, brain and muscle.

**Figure 2.**
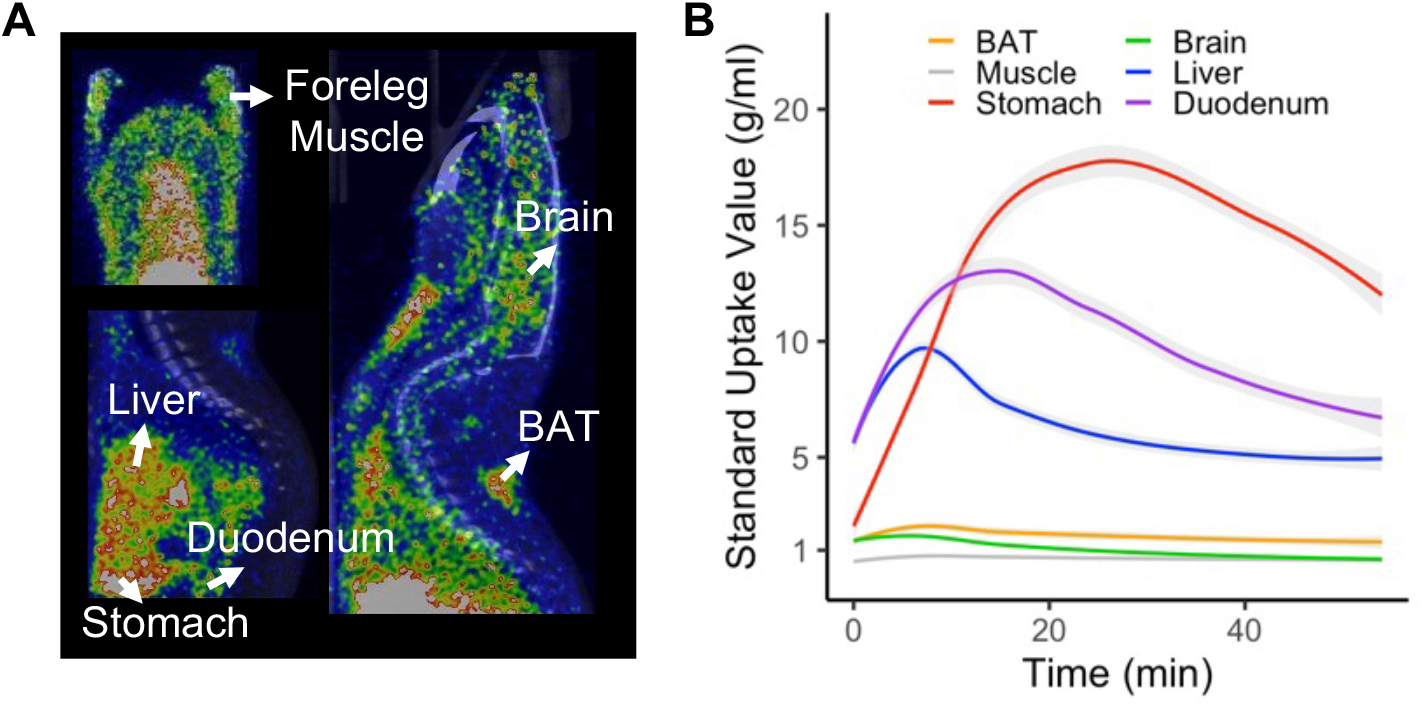
[^11^C]carfentanil standardized update values (SUVs) in different tissues. **A.** PET-CT fusion image of a rat, with regions of interest labelled. **B.** Regional time-activity curves of the SUVs across scans. Shaded areas represent 95% Confidence Interval (CI).

### Tissue-specific decomposition of [^11^C]carfentanil

To rule out potential non-specific binding of radioactive metabolites, we studied the metabolic profiles of [^11^C]carfentanil in different tissues. Analysis of blood samples demonstrated a continuous decomposition of the radiotracer (**Fig. 3A&B**). Measures in seven rats (**Fig. 3B**) confirmed that, at 20 minutes after radiotracer injection, the SUVs in liver and duodenum were vastly driven by uptake of radioactive metabolites. Therefore, [^11^C]carfentanil PET is not an optimal tool to study the MOR availability in these tissues. Comparable profiles of [^11^C]carfentanil radiometabolism were found in BAT, muscle and stomach, with [^11^C]carfentanil contributing over 50% of the total radioactivity at 20 minutes after radiotracer injection. The brain had the highest composition of intact [^11^C]carfentanil.

**Figure 3.**
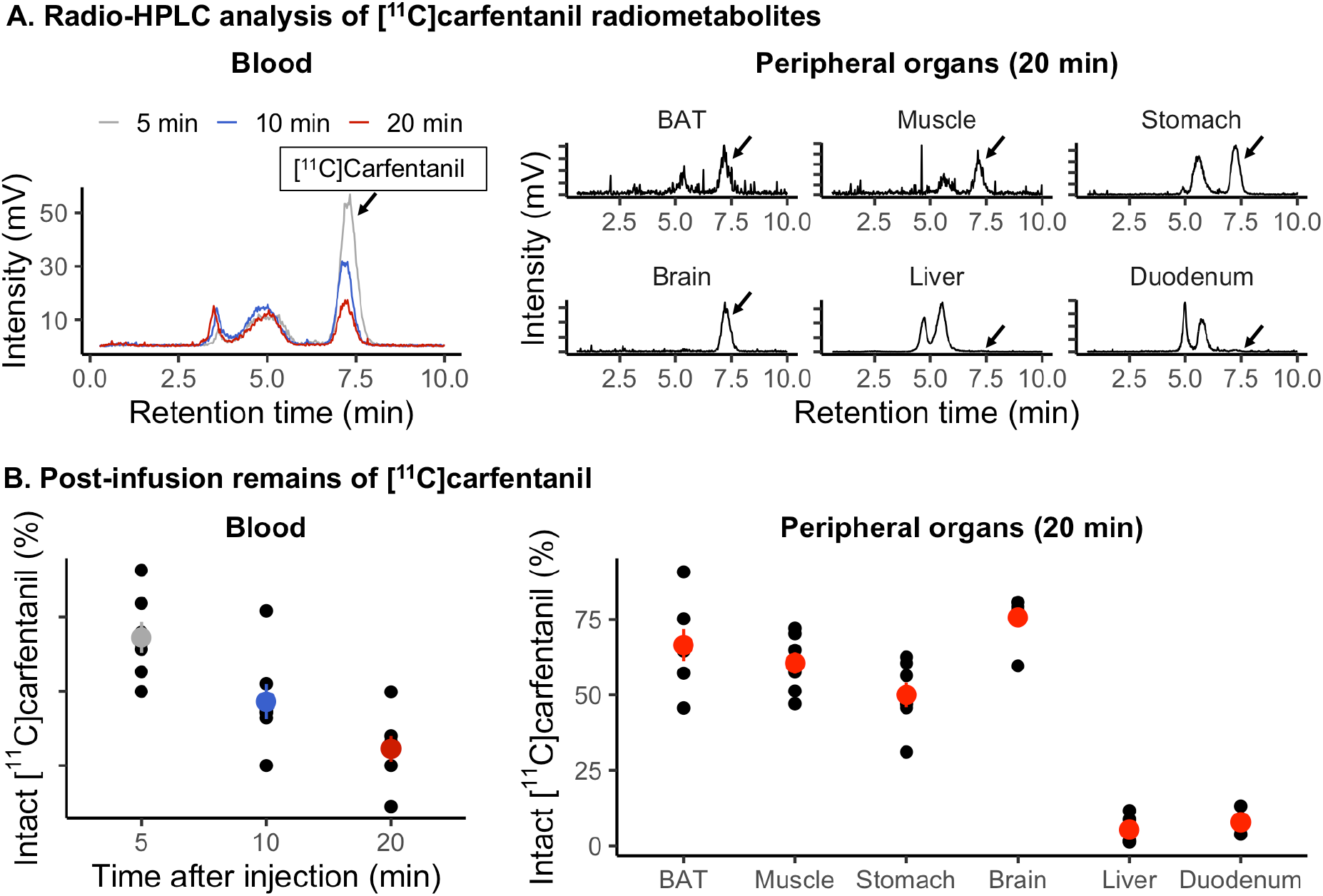
Metabolic decomposition of [^11^C]carfentanil in distinct rat tissues after intravenous injection of radiotracer. **A.** HPLC measures of regional decompositions of [^11^C]carfentanil. Measures of one rat were used for demonstration. **B.** Regional remains of [^11^C]carfentanil at different time points. Seven rats were studied. Error bars represent the 95% CI.

In the stomach, around 50% [^11^C]carfentanil remained at 20 min after tracer injection, and SUVs showed much higher levels of MOR availability. However, stomach was not scanned in some images, and therefore stomach data was not further analyzed.

### [^11^C]carfentanil uptake as a function of photoperiod

TACs of [^11^C]carfentanil SUVs across scans in BAT and muscle were plotted aligning to the length of photoperiods (**Fig. 4**). There was a pattern showing that, areas under the TACs in BAT were lower toward longer photoperiod. BAT had higher radiotracer uptake than muscle in all scans.

**Figure 4.**
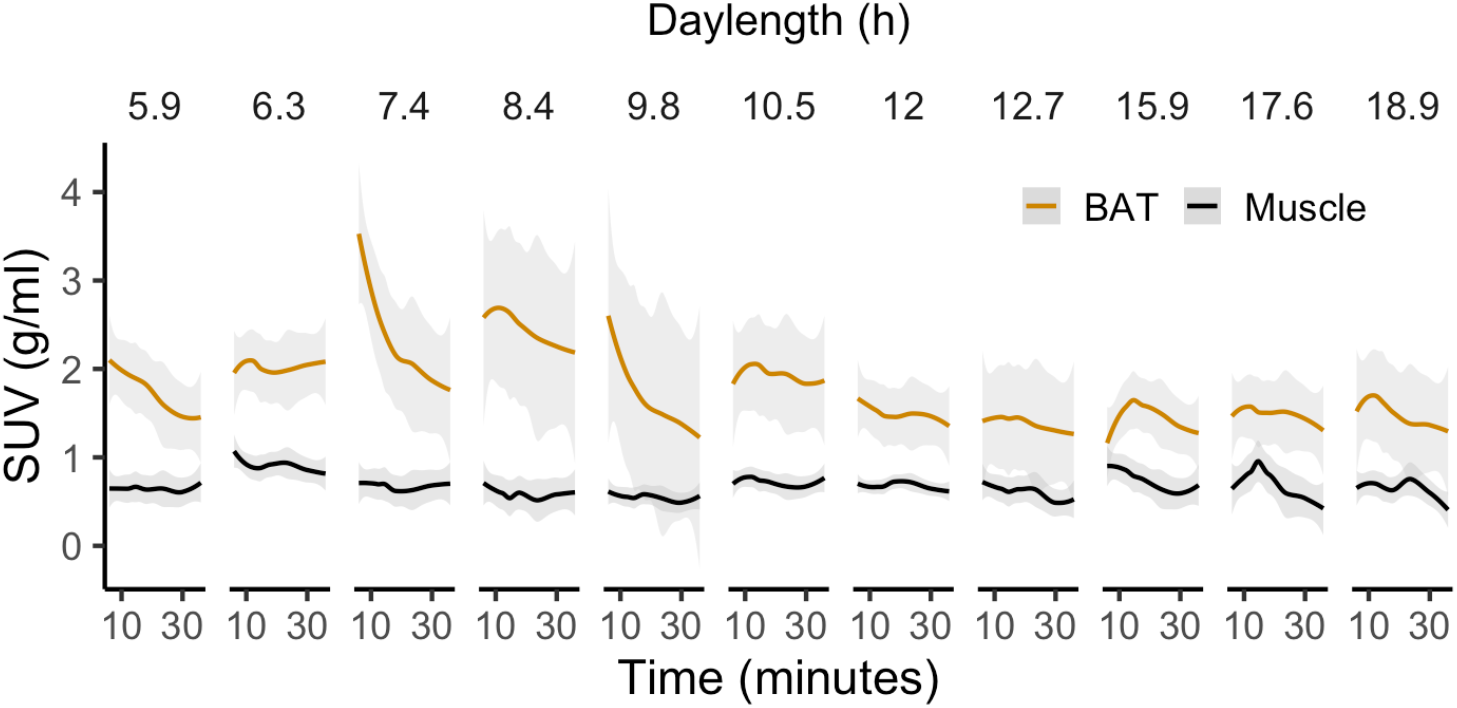
Time-activity curves (TACs) in BAT and muscle as a function of photoperiod. Time range between 5-40 minutes for the TACs are shown. Shaded areas represent 95% CI.

### Effect of photoperiod on BAT MOR availability

While the SUVs (**Fig. 2B**) and the metabolic profiles of [^11^C]carfentanil (**Fig. 3A**) were comparable in BAT and muscle, BAT-to-muscle SUV ratios was firstly used to estimate the MOR availability in BAT.

In the statistical analysis of MOR availability, firstly, both the experimental group (n = 3) and control group (n = 9) rats were involved. We compared three statistical models: i) Model 1 with Scanner Type as the only fixed effect factor, ii) Model 2 with Scanner Type and Photoperiod as fixed effect factors, and iii) Model 3 with Scanner Type, Photoperiod and Age as fixed effect factors. In all these mixed-effect models, Rats were used as a random factor. Results showed that Model 2 was the most optimal model with the lowest Akaike information criterion (AIC) values (Model 1: 71.4; Model 2: 67.9; Model 3: 69.9) and the largest marginal R^2^ (Model 1: 0.09; Model 2: 0.18; Model 3: 0.17). In Model 2, Photoperiod had significant effect on MOR availability in BAT (β = −0.04, 95% CI [−0.08, −0.01]). In Model 3, age had no effect (β = −0.0002, 95% CI [−0.006, 0.006]) on MOR availability, and therefore Age as a factor was dropped from following analysis.

Statistical analysis was also done by including only the experimental group rats (n = 9), which were under continuously changing photoperiod. MOR availability at BAT were analyzed using fixed-effect factors including Scanner Type and Photoperiod, and Rats as random factor. Similarly, increasing daylength led to reduced peripheral MOR binding in BAT (β = −0.04, 95% CI [−0.07, −0.01]). Plot of SUV ratios (**Fig. 5A)** demonstrated that photoperiod had an inversed association with MOR availability in BAT.

**Figure 5.**
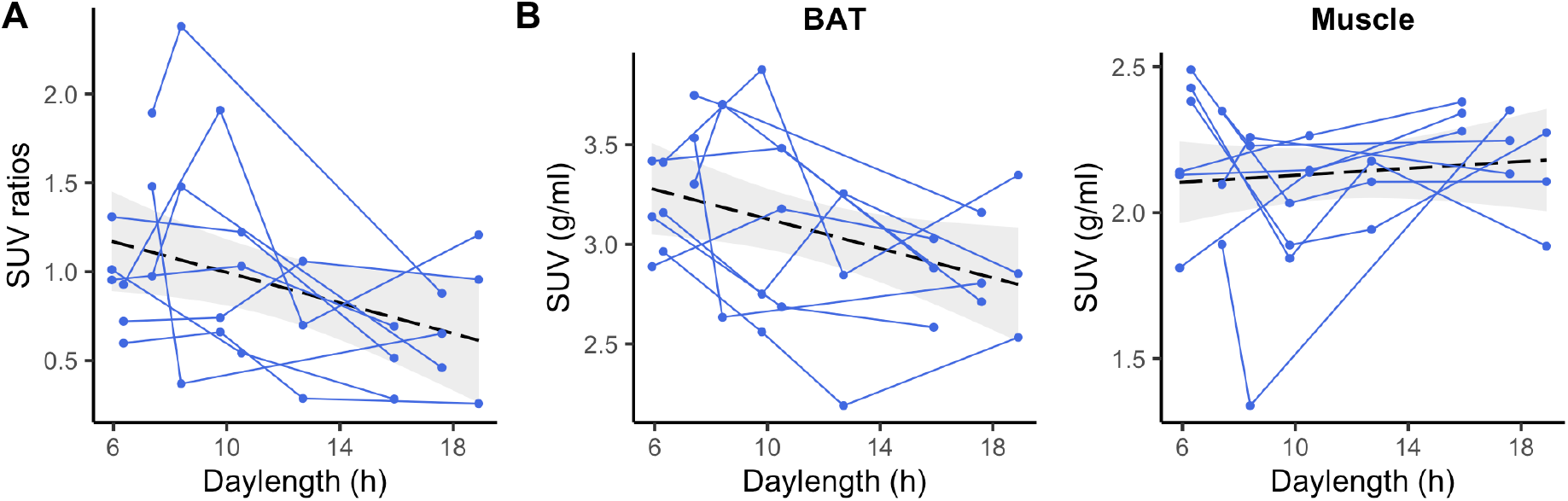
Plot of SUV ratios and SUVs demonstrates the effect of photoperiod on MOR availability. **A.** Plot of SUV ratios between BAT and muscle. **B.** Plots of SUVs separately for BAT and Muscle. Individual animals are shown as separate lines. Black dashed line shows the least-squares (LS) regression line for the linear model (y ~ x) predicting SUV ratios/SUVs across the whole sample. Shaded area shows the 95% CI for the LS curve.

### Statistical analysis of BAT and muscle SUVs

Additionally, we analyzed the [^11^C]carfentanil SUVs separately for BAT and Muscle. Areas under TACs (5 - 20 min) were used as indices of the MOR availability. When involving both the control and experimental group, increasing photoperiod led to lower MOR availability in BAT (β = −0.037, 95% CI [−0.07, −0.01]) but not in muscle (β = 0.005, 95% CI [−0.01, 0.02]). When involving only the experimental group, similar results were found with increasing photoperiod associated with lower MOR availability in BAT (β = −0.04, 95% CI [−0.06, −0.01]) but not in muscle (β = 0.006, 95% CI [−0.01, 0.02]). Plots for SUVs (**Fig. 5B**) showed that photoperiod had an inversed association with MOR availability in BAT but not muscle.

## Discussion

Our main finding is that photoperiod modulates MOR availability in BAT, with longer photoperiod causally leading to lower MOR availability. The lowered MOR availability in BAT during bright seasons may suggest lowered innervation of BAT thermogenesis, which aligns with the reduced BAT thermogenesis during summer (McElroy and Wade, 1986; Au- Yong et al., 2009). The neurotransmitter signaling underlying functional modulations of BAT metabolism are poorly understood. The current data suggest that MOR signaling is an important neurochemical mediator for BAT activity. Together with our previous findings on photoperiod-tuned central MOR system (Sun et al., 2021), data reveal that both central and peripheral MOR signaling pathways possess seasonal patterns. The study also highlights the applicability of [^11^C]carfentanil PET in studying *in vivo* peripheral MOR signaling.

Exact mechanism of photoperiod modulated MOR availability in BAT remains elusive. The MOR signaling pathway may carry communicative signals within the gut-brain-BAT axis. MORs are found throughout the central and enteric nervous system (Sternini et al., 2004) with interactive effects on gastrointestinal motility, possibly regulating feeding behavior (Janssen et al., 2011). Also, activation of central MOR signaling with fentanyl is found to elicit BAT thermogenesis (Cao and Morrison, 2005; Cao et al., 2010). While previous studies mostly focus on the efferent modulatory effect of brain to gut or BAT, afferent feedback of BAT activation to brain functions have been also proposed (Li et al., 2018; Laurila et al., 2021), despite of the lack of MOR-related evidence. The current study shows that photoperiod not only modulates the central MOR availability but also the receptor availability in BAT. This suggests a possible novel communicative approach between BAT and brain. Based on these assumptions, lowered MOR availability in BAT may suggest reduced active communication between brain and BAT during bright seasons.

Our finding that photoperiod leads to decreased MOR availability is also possibly caused by increased peripheral opioid release, which competes for MOR binding with the radiotracer, during bright seasons. Peripheral opioid signaling is elevated during inflammatory reactions due to the secretion of endogenous opioid by immune cells (Hassan et al., 1993; Kraus et al., 2001; Stein et al., 2001). Inflammatory response in the BAT leads to reduced capacity of energy expenditure and glucose uptake (Omran and Christian, 2020), which is in line with the subsequently reduced thermogenesis in BAT during long days (McElroy and Wade, 1986; Au- Yong et al., 2009). However, this possible pathway lacks direct support from the current study. The central MOR signaling contributing to hedonic or “liking” responses in the brain (Berridge, 2009) are related to the brain food-reward response (Nummenmaa et al., 2018). Obese subjects show lowered brain MOR levels and bariatric surgery normalizes their levels (Karlsson et al., 2016), highlighting the role of central MOR in feeding behavior and weight gain. Recently, the MOR antagonist drug naltrexone, as part of the *Contrave* complex, has been approved by the U.S. Food and Drug Administration for treating obesity. While *Contrave* shows effectiveness in reducing weight in a randomized clinical trial (Apovian et al., 2013), the exact mechanism is still unclear. The other component of the *Contrave* complex is bupropion which is a dopamine and norepinephrine reuptake inhibitor, which together with naltrexone is considered to modulate food cravings through an effect on the reward pathways. While central MOR signaling is a most probable effector for this drug effectiveness regarding for instance food intake habits, peripheral MOR signaling functions should not be neglected. So far, no previous studies address this aspect.

Immunofluorescence staining data reveal expression of MOR in both central and peripheral tissues, affirming the [^11^C]carfentanil PET findings in the current study, as well as a previous whole-body human [^11^C]carfentanil PET study (Newberg et al., 2009). While, extensive studies have correlated the levels of brain MOR signaling with socioemotional functions, understanding of the potential roles of peripheral MOR signalling in cognitive functions and behaviour, however, is still at the infant stage. Whether and how peripheral MOR signal compensates central MOR activity in regulating human reward-related responses and trait-level behaviour requires in-depth investigation.

The current study shows that [^11^C]carfentanil PET can be used to measure MOR availability in peripheral tissues where the accumulation of radioactive metabolites of [^11^C]carfentanil is not significant. Although gastric MOR functions are not covered by this study, our data encourage future studies to investigate how *in vivo* MOR signaling may affect gastric function and eating behavior. Further, the weight-based doses of [^11^C]carfentanil in imaged rats are comparable to those used in routine clinical imaging studies (Hirvonen et al., 2009), and therefore [^11^C]carfentanil PET could be also used to study peripheral MOR functions in humans.

## Limitations

While we find that photoperiod modulates both BAT and brain MOR signaling, the patterns are largely distinct, i.e., the linearly reverse vs. inverted-U functional relationships. Brain and BAT have comparable amount of MOR density (see **Fig. 1&2**), but much higher amount of MOR distribution is found in other peripheral tissues. For instance, the stomach has much higher MOR availability compared to brain and BAT. We are not aware of how these relative densities affect the endogenous opioid signaling, subsequently influencing feeding behavior and weight gain. Furthermore, the current study does not involve MRI measures to evaluate the BAT mass, and therefore we cannot rule out potential effect of photoperiod on BAT volume changes (McElroy and Wade, 1986) which affects [^11^C]carfentanil uptake. Also, although the current study demonstrates that photoperiod is negatively correlated with MOR availability in BAT, it cannot answer the question whether changed MOR availability is causally associated with BAT metabolic levels and thermogenesis.

## Conclusion

Our data have shown that photoperiod modulates the BAT MOR availability. This extends our previous finding that photoperiod shapes brain MOR availability, to reveal a shared neurotransmitter signaling pathway between brain and body in reacting to seasonal rhythms. Seasonal affective disorders are characterized with winter blue and overeating. Considering the important role of MOR signaling in emotion, eating behavior, thermogenesis, as well as the important role of BAT in weight gain and motivation of eating (Li et al., 2018; Laurila et al., 2021), the study further supports that the MOR signaling pathway may be underlying the seasonal affective changes. Finally, this study justifies the feasibility of [^11^C]carfentanil PET in studying peripheral MOR signaling.

